# Yatakemycin biosynthesis requires two deoxyribonucleases for toxin self-resistance

**DOI:** 10.1101/2024.09.16.613364

**Authors:** Jonathan Dorival, Hua Yuan, Allison S. Walker, Gong-Li Tang, Brandt F. Eichman

**Affiliations:** Department of Biological Sciences, Vanderbilt University, Nashville, Tennessee, USA; State Key Laboratory of Chemical Biology, Shanghai Institute of Organic Chemistry, University of Chinese Academy of Sciences, Chinese Academy of Sciences, Shanghai 200032, China; College of Life Sciences, Shanghai Normal University, Shanghai 201418, China; Department of Chemistry, Vanderbilt University, Nashville, Tennessee, USA; School of Chemistry and Materials Science, Hangzhou Institute for Advanced Study, University of Chinese Academy of Sciences, 1 Sub-lane Xiangshan, Hangzhou 310024, China; Department of Biochemistry, Vanderbilt University School of Medicine, Nashville, Tennessee, USA

## Abstract

The highly active natural product yatakemycin (YTM) from *Streptomyces* sp. TP-A0356 is a potent DNA damaging agent with antimicrobial and antitumor properties. The YTM biosynthesis gene cluster (*ytk*) contains several toxin self-resistance genes. Of these, *ytkR2* encodes a DNA glycosylase that is important for YTM production and host survival by excising lethal YTM-adenine lesions from the genome, presumably initiating a base excision repair (BER) pathway. However, the genes involved in repair of the resulting apurinic/apyrimidinic (AP) site as the second BER step have not been identified. Here, we show that *ytkR4* and *ytkR5* are essential for YTM production and encode deoxyribonucleases related to other known DNA repair nucleases. Purified YtkR4 and YtkR5 exhibit AP endonuclease activity specific for YtkR2-generated AP sites, providing a basis for BER of the toxic AP intermediate produced from YTM-adenine excision and consistent with co-evolution of *ytkR2, ytkR4*, and *ytkR5*. YtkR4 and YtkR5 also exhibit 3′-5′ exonuclease activity with differing substrate specificities. The YtkR5 exonuclease is capable of digesting through a YTM-DNA lesion and may represent an alternative repair mechanism to BER. We also show that *ytkR4* and *ytkR5* homologs are often clustered together in putative gene clusters related to natural product production, consistent with non-redundant roles in repair of other DNA adducts derived from genotoxic natural products.

## Introduction

Some plant and microbial natural products are toxic by virtue of their ability to chemically modify DNA. Genotoxic compounds are diverse in chemical structure and generate an array of covalent and non-covalent DNA adducts that inhibit DNA processing and lead to cell death or disease ^1–7^. The high cytotoxicity of DNA damaging natural products can be harnessed to develop antimicrobial and anticancer drugs such as actinomycin D, daunomycin, mitomycin C, and calicheamicins ^1^. Cells have evolved several conserved pathways to detect and repair different types of DNA damage as a means of survival ^2, 8–10^. For example, bulky, helix-distorting adducts are typically removed by nucleotide excision repair (NER), whereas smaller lesions are repaired by direct reversal or base excision repair (BER) pathways.

Yatakemycin (YTM, **1**) is produced by *Streptomyces* sp. TP-A0356 and belongs to the spirocyclopropylcyclohexadienone (SCPCHD) family of genotoxic natural products that includes duocarmycin A, and duocarmycin SA, CC-1065, and gilvusmycin (Fig. 1A) ^11–15^. These compounds preferentially bind the minor groove of AT-rich sequences and alkylate the N3-position of adenine through ring opening of their cyclopropyl groups (Fig. 1B) ^16–19^. In addition to covalent adducts, SCPCHD compounds also form a network of non-covalent interactions with both DNA strands that stabilizes the duplex and effectively creates a non-covalent interstrand DNA crosslink that poses a challenge to excision repair pathways ^20–24^. Consequently, SCPCHD-DNA adducts are potent blocks to DNA replication and exhibit antibiotic, antifungal, and antitumor properties ^11–15, 25–28^. YTM is unique in that its cyclopropyl ring resides in the middle subunit, which forms a “sandwiched” structure with an enhanced rate of DNA alkylation, making YTM is the most potent SCPCHD member with an IC_50_ of 3-5 pM against the L1210 cell line ^17^.

**Figure 1.**
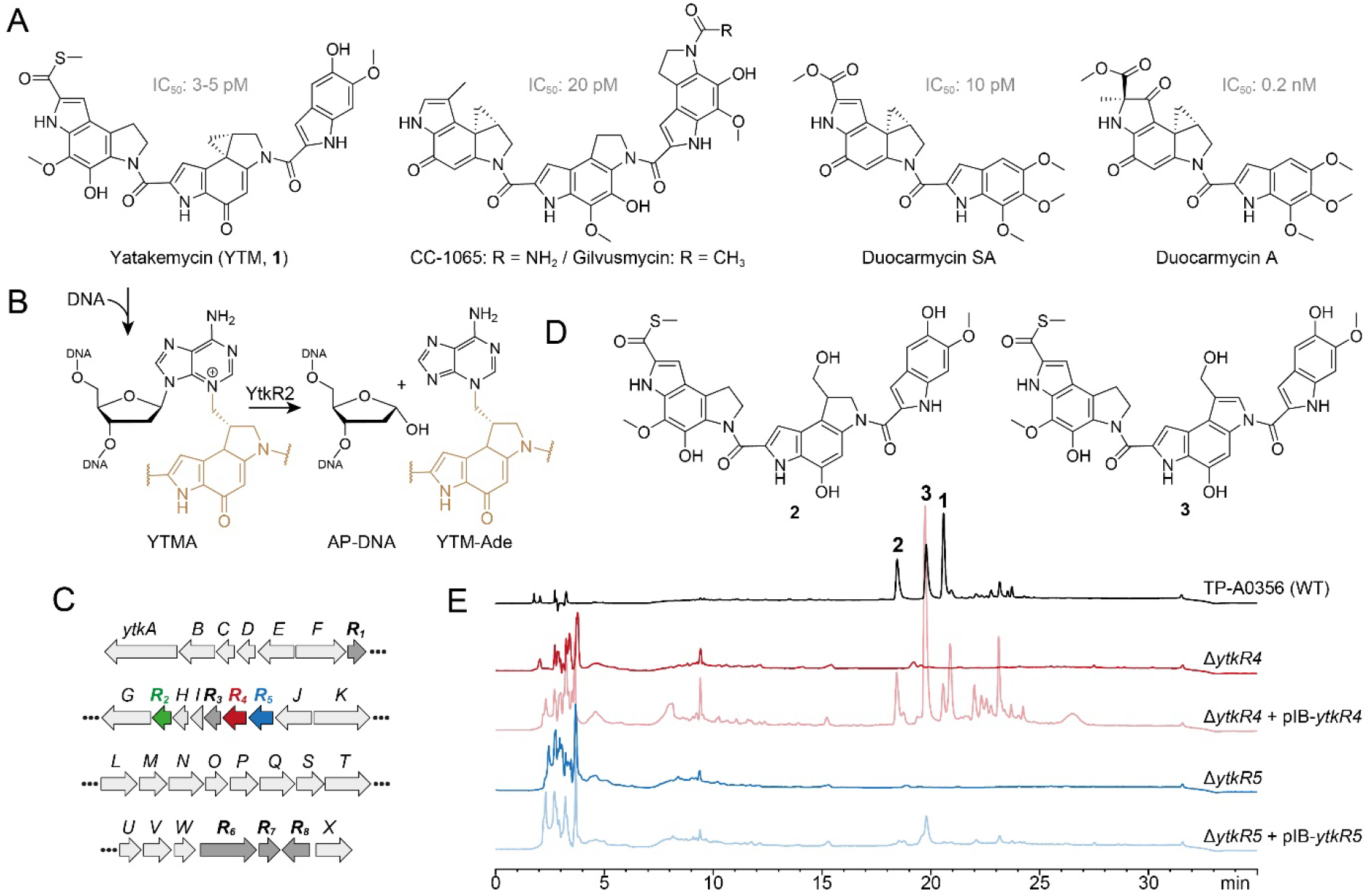
Effects of ytkR4 and ytkR5 on yatakemycin biosynthesis. (A) Structures of yatakemycin (YTM, **1**), CC-1065, gilvusmycin, and duocarmycins. (B) YtkR2-catalyzed hydrolysis of a YTMA-DNA lesion yields an AP site in the DNA and free YTM-Ade. The YTM moiety is colored tan. (C) YTM biosynthetic gene cluster. Resistance (*ytkR*) genes are labeled bold and highlighted in color or dark grey. (D) YTM hydrolysis products, **2** and **3**. (E) HPLC traces of fermentation products extracted from the wild-type TP-A0356 strain and Δ*ytkR4* and Δ*ytkR5* mutants. Each mutant was analyzed alone and complemented with the corresponding native *ytkR4* or *ytkR5* gene (pIB139 vector). The injected concentration of the extracted fermentation products from TP-A0356 is 1/20 that of the other strains. The detection wavelength used was 383 nm.

Toxin-producing microorganisms have self-resistance mechanisms for survival and, in the case of genotoxins, to protect the integrity of the genome ^29–37^. These self-resistance mechanisms are often embedded within and co-evolve with the biosynthesis gene cluster (BGC). Biosynthetic studies revealed that the YTM producer encodes multiple resistance/regulation genes, *ytkR1*-*ytkR8*, in its BGC (Fig. 1C) ^26^. The *ytkR6* gene is homologous to drug-resistance transporters and was proposed to function as an efflux pump ^26^. Three genes—*ytkR1, ytkR7*, and *ytkR8*—encode GyrI-like small molecule binding proteins; YtkR7 inactivates YTM by catalyzing the hydrolysis of the cyclopropyl warhead to form products **2** and **3** (Fig. 1D), and YtkR1 and YtkR8 have been proposed to sequester YTM or its intermediates and to regulate expression of the cluster, respectively ^38^.

YtkR2 confers self-resistance against YTM by functioning as a DNA glycosylase that hydrolyzes YTM-adenosine (YTMA) adducts to yield free YTM-adenine (YTM-Ade) (Fig. 1B) ^39^. DNA glycosylases initiate the BER pathway by hydrolyzing the N-glycosidic bond of the aberrant nucleotide to generate an apurinic/apyrimidinic (AP, or abasic) site. The AP site is a reactive intermediate that is removed in the subsequent steps of BER, whereby an AP endonuclease hydrolyzes the phosphodiester bond on the 5′ side of the AP site to generate a 3′-hydroxyl for gap filling synthesis by a DNA polymerase, followed by nick sealing by DNA ligase ^40^. YtkR2 is a member of the AlkD family of DNA glycosylases that are unique in their ability to excise bulky DNA adducts ^20–22, 39, 41–46^. Consistent with its co-evolution within the YTM cluster, YtkR2 confers a greater cellular resistance against YTM than does *Bacillus cereus* AlkD ^22^. This enhanced resistance is not the result of substrate specificity, but rather from a low affinity of YtkR2 for its AP site product, which presumably allows for enzymatic removal of the AP site ^20, 22^. How the AP site is repaired in *Streptomyces* sp. TP-A0356, however, has been unclear.

Here, we show genetically and biochemically that *ytkR4* and *ytkR5* encoded in the YTM BGC (Fig. 1C) are resistance genes that encode multifunctional deoxyribonucleases (DNases) capable of processing YTMA-DNA adducts. YtkR4 is homologous to the TatD family of DNases, and YtkR5 is a putative member of the xylose isomerase-like TIM barrel domain family found in the bacterial AP endonuclease, Endonuclease (Endo) IV ^26, 39^. Consistent with these annotations, we found that YtkR4 and YtkR5 are 3′-5′ exonucleases with a preference for single-(ss) and double-stranded (ds) DNA substrates, respectively, and that YtkR5 can degrade DNA containing a YTMA lesion. In addition, both enzymes exhibit AP endonuclease activity toward the toxic AP product of YTMA hydrolysis by YtkR2. We also found through bioinformatic analysis that *ytkR4* and *ytkR5* homologs in other bacteria are often located together within putative BGCs or other gene clusters related to natural product production, suggesting they play multiple, non-overlapping roles in repair of genotoxin DNA adducts.

## Results and discussion

### YtkR4 and YtkR5 are essential for yatakemycin biosynthesis

To investigate the effects of *ytkR4* and *ytkR5* on YTM biosynthesis, we generated Δ*ytkR4* and Δ*ytkR5* mutants of the

YTM producing strain *Streptomyces* sp. TP-A0356 by constructing an in-frame deletion for each gene through homologous recombination (Supplementary Fig. S1). We then carried out fermentation analyses and monitored YTM production by HPLC. Both mutants lost the ability to produce YTM and its two hydrolyzed products, **2** and **3** (Fig. 1E). To verify the loss of YTM production was related to YtkR4 and YtkR5 activities, we introduced a plasmid expressing the native gene into the corresponding mutant. Complementation partially restored production of YTM and/or its two hydrolyzed products in each mutant (Fig. 1E). Thus, these genetic results indicate that the *ytkR4* and *ytkR5* genes are each essential for YTM biosynthesis.

### YtkR4 and YtkR5 exhibit exonuclease and AP endonuclease activities

YtkR4 and YtkR5 are distant homologs of TatD and EndoIV deoxyribonucleases, respectively (Supplemental Figs. S2 and S3) ^26, 39^. Both TatD and EndoIV exhibit AP endonuclease and 3′-5′ exonuclease activities (Fig. 2A) ^47–52^. We therefore tested nuclease activities of purified YtkR4 and YtkR5 proteins on 5′-FAM labeled ssDNA and dsDNA substrates containing either no modification or a centrally located tetrahydrofuran (THF) abasic site analog (Fig. 2B). THF is more stable than a natural AP site owing to the lack of the hydroxyl group at deoxyribose C1′, and is a substrate for other AP endonucleases including TatD and EndoIV ^47, 53^. In the presence of Mg^2+^ cofactor, YtkR4 and YtkR5 generated a ladder of products on both ssDNA and dsDNA, indicative of 3′-5′ exonuclease activity, compared to no-enzyme controls (Fig. 2C). YtkR4 showed greater exonuclease activity on ssDNA than on dsDNA, similar to the substrate specificity of human and *E. coli* TatDs ^47, 49^. In contrast, YtkR5 exhibited a preference for dsDNA, consistent with its homology to EndoIV, which also has greater activity for dsDNA ^52^. In addition to exonuclease activity, both enzymes displayed AP endonuclease activity on the dsDNA THF substrate, as evidenced by the accumulation of a band corresponding to specific cleavage at the THF (Fig. 2C). YtkR5, but not YtkR4, showed a robust exonuclease degradation of the product of the AP endonuclease reaction, consistent with its preference for exonucleolytic activity in dsDNA. Neither enzyme exhibited AP endonuclease activity in ssDNA. THF inhibited the ssDNA exonuclease activity of YtkR4, as we observed an accumulation of a band corresponding to an oligonucleotide one nucleotide longer than the AP endo cleavage product on the THF-ssDNA but not the unmodified ssDNA substrate, indicating that YtkR4 pauses after cleaving the nucleotide immediately 3′ to the THF. We previously observed the same pausing behavior in response to THF from the human and *E. coli* TatD enzymes ^47^. We also examined the effect of adding both YtkR4 and YtkR5 to the reactions and found that while the incision products represented a sum of the individual reactions, exonuclease degradation of the AP site product was modestly faster (Supplementary Fig. S4).

**Figure 2.**
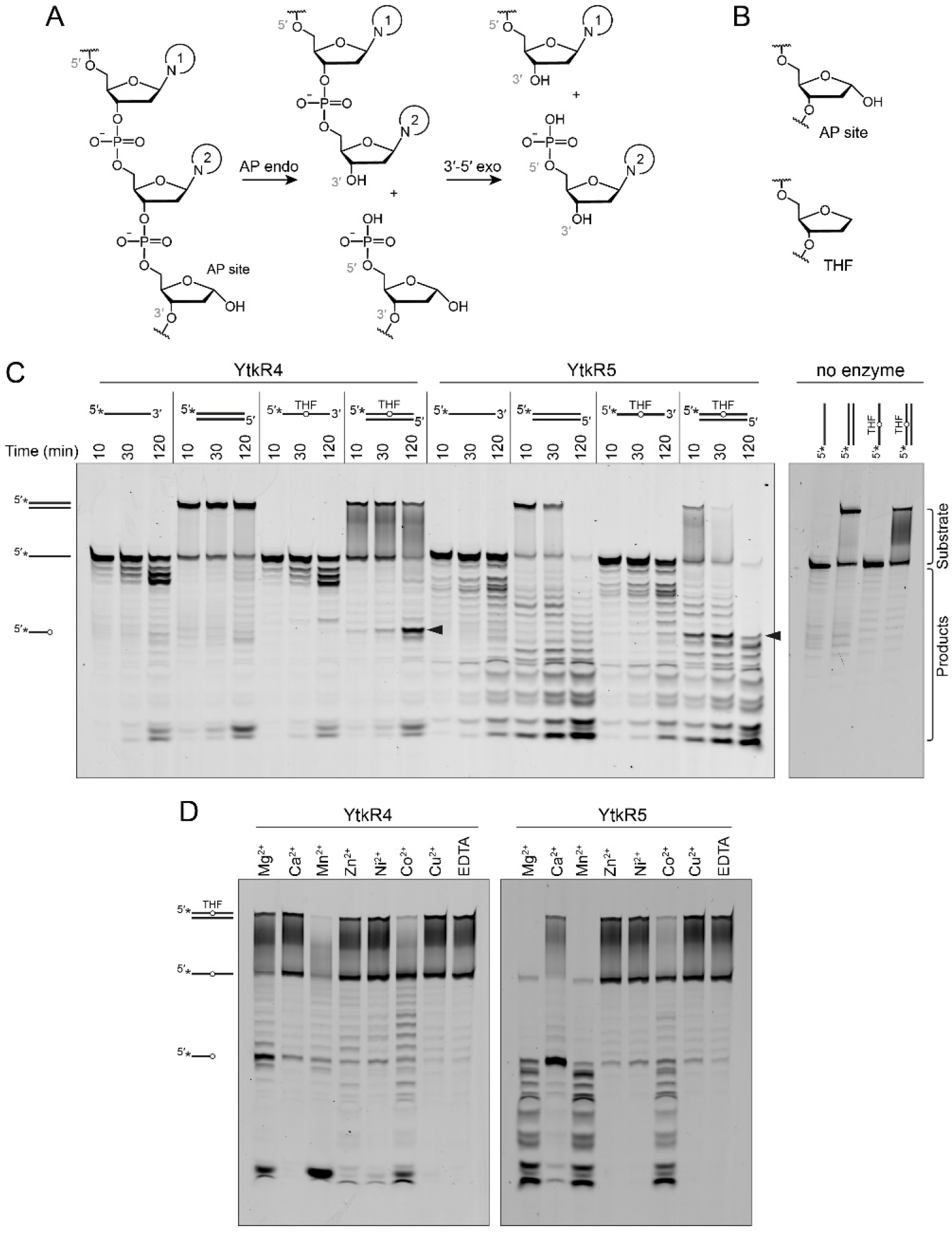
YtkR4 and YtkR5 exhibit 3′-5′ exonuclease and AP endonuclease activity. (A) AP endonuclease and 3′-5′ exonuclease chemical reactions. Nucleobases are labeled 1 and 2 for identification purposes. (B) Chemical structure of the tetrahydrofuran (THF) abasic analog compared to a natural AP site. (C) Denaturing PAGE of 5′-FAM-labeled substrates incubated with YtkR4 or YtkR5 for the indicated times or with buffer alone (no enzyme) for 120 min. Asterisks (*) in the substrate schematics denote the location of the FAM label. Bands corresponding to substrates and products are indicated to the right of the gel. Black triangles designate the bands resulting from AP endonuclease activity. Duplex DNA persists on the gel because of the high GC content (Supplementary Table S3). (D) Metal dependence for nuclease activity on a 5′-FAM-labeled double-stranded THF-DNA substrate. Reactions were carried out for 2 hr (YtkR4) or 90 min (YtkR5) and contained 10 µM protein, 100 nM DNA, and either 10 mM MgCl_2_, 5 mM CaCl_2_, 0.05 mM ZnCl_2_, 3 mM MnCl_2_, 1 mM NiCl_2_, 1 mM CuCl_2_, or 10 mM EDTA.

Since TatD and EndoIV utilize metal cofactors, we examined the metal dependence of YtkR4 and YtkR5 (Fig. 2D). We tested the activity of each protein against seven different divalent cations using the THF-containing dsDNA substrate so that we could monitor both AP endo and exonuclease activities. YktR4 exhibited the most exonuclease activity in the presence of Mg^2+^, Mn^2+^, and Co^2+^, weak activity with Ca^2+^, Zn^2+^, and Ni^2+^, and no activity with Cu^2+^. AP endonuclease activity of YtkR4 was greatest in presence of Mg^2+^, and a low level of activity was observed with Ca^2+^. YktR5 exhibited both AP endonuclease and exonuclease activity in presence of Mg^2+^, Mn^2+^, and to a lesser extent Co^2+^. In the presence of Ca^2+^, YtkR5 displayed robust AP endonuclease activity and reduced exonuclease activity. Thus, whereas YtkR4 exhibited the same metal dependence as the TatD enzymes ^47^, the metal dependence of YtkR5 was markedly different than EndoIV. While EndoIV uses Zn^2+^ as a preferred metal for activity and remains active in presence of EDTA ^52^, YtkR5 exhibited the most nuclease activity in the presence of Mg^2+^, Mn^2+^ and Ca^2+^ and very low activity in the presence of Zn^2+^ and EDTA. This indicates that YtkR5 may have evolved differently compared to EndoIV.

The metal preferences can be explained by similarities and differences in metal binding residues in the active sites of these enzymes (Supplementary Fig. S2 and S3). YtkR4’s substrate specificity and metal dependence are reminiscent of the TatD enzymes ^47, 49^. Despite only 20% sequence identity overall, the TatD metal binding residues are largely conserved in YtkR4, with only two out of six exceptions (Supplementary Fig. S2). YtkR4 Arg168 and Gly241 align with conserved TatD His and Asp residues, respectively. Although these substitutions would not coordinate the metals, they are not expected to alter the metal preference because they are each one of four that bind a separate metal ^47^. YtkR5, on the other hand, has a markedly different metal dependence than EndoIV despite the similarity in preference for dsDNA ^52^. Several EndoIV residues known to bind metals are different or absent in YtkR5, which likely affect the nature or number of metals bound in the active site (Supplementary Fig. S3). For example, Zn_A_ ^2+^ and Zn_B_ ^2+^ binding residues Asp229 and His216 in EndoIV align with His218 and Gln209 in YtkR5. EndoIV His69 and His109, which bind the third Zn^2+^ ion, align with Asp77 and Ala117 in YtkR5, such that only Glu155 would be involved in the binding of the third metal.

### YtkR4 and YtkR5 AP endonucleases have a modest preference for the YtkR2 product

YtkR4 and YtkR5 presumably evolved with the *ytk* gene cluster, and thus we tested the hypothesis that these nucleases would show a preference for the products of YtkR2 cleavage of YTMA lesions (i.e., AP-DNA + YTM-Ade) (Fig. 1B). We examined their AP endonuclease activities on substrates containing natural abasic sites generated either by uracil DNA glycosylase (UDG) excision of deoxyuracil or by YtkR2 excision of YTMA, as compared to the THF-containing substrate (Fig. 3A). The total (AP endo + exo) nuclease activity of YtkR4 was similar among the three substrates, as judged by the rates of product accumulation, whereas YtkR5 exhibited slightly higher overall nuclease activity for THF or UDG-derived AP sites (Fig. 3B). However, both enzymes showed slightly higher AP endo activity toward the YTMA-derived AP site (Fig. 3C). In the case of YtkR4, the incision product of the YTMA-derived AP site accumulated faster and to a greater extent than those of the UDG-derived or THF sites. In contrast, the products of YtkR5 AP endo activity from YTMA-derived AP sites accumulated to a greater extent, albeit slower, than those of the UDG-derived or THF AP sites (Fig. 3C). Together, these results indicate that the AP endonuclease activities of both YtkR4 and YtkR5 are specific for AP sites derived from YtkR2 excision of YTMA. Because the AP sites produced from YtkR2 and UDG are identical, the specificity is likely the result of the presence of the excised YTM-Ade nucleobase remaining non-covalently bound in the DNA. YTM makes intimate interactions with both DNA strands ^20^ and thus we would not expect YTM-Ade to dissociate as readily as uracil. We previously showed that the affinity of YtkR2 for its AP site product is greater in the presence of the excised YTM-Ade adduct ^22^, and thus the specificity of YtkR4 and YtkR5 for YtkR2-derived AP sites may also be attributed to a potential interaction with YtkR2 prior to its dissociation from the AP site. In either scenario, either the excised YTM-Ade adduct or YtkR2 could help guide the nuclease to the site of damage through a direct interaction but would also require dissociation to enable the nuclease to fully access the AP site for incision.

**Figure 3.**
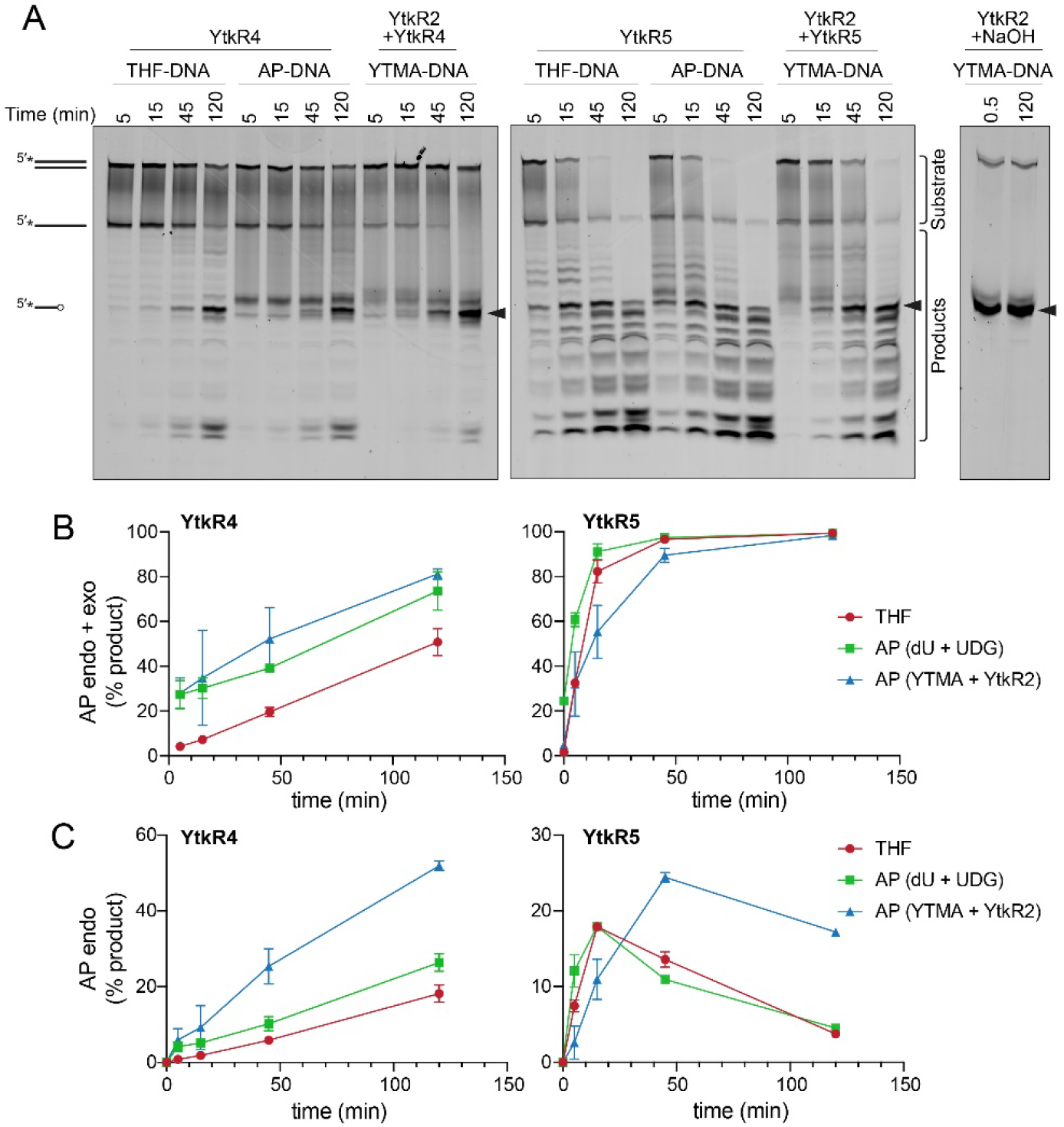
YtkR4 and YtkR5 have greater AP endonuclease activities on abasic sites generated by YtkR2 excision of YTMA. (A) Nuclease activities of YtkR4 and YtkR5 on dsDNA substrates containing either THF (THF-DNA), an AP site generated by UDG excision of deoxyuracil (AP-DNA), or the product of YtkR2 excision of YTMA (YTMA-DNA). The far-right panel shows non-enzymatic (NaOH) cleavage of the AP site generated from the YtkR2/YTMA-DNA glycosylase reaction. Bands quantified as substrates and products are indicated. Black triangles designate the bands resulting from AP endonuclease activity. (B) Quantification of both AP endo and exonuclease products. Data are represented as the mean ± SD (n = 2). (C) Quantification of AP endonuclease products. Mean ± SD (n = 2).

### Exonuclease activity of YtkR5 is not inhibited by YTMA lesions

Because of the possibility that YtkR4 and YtkR5 would encounter a YTMA lesion in DNA prior to its excision by YtkR2, we examined whether this adduct inhibits the exonuclease activities of YtkR4 and YtkR5. We incubated each enzyme with a dsDNA substrate containing a centrally located YTMA lesion and compared the kinetics of exonuclease activity against the same substrate containing no modification (Fig. 4). Compared to its activity on unmodified DNA, YtkR4 did not produce excision products 5′ to the lesion, indicating that it is unable to digest DNA beyond YTMA (Fig. 4A). Interestingly, YtkR4 experienced a slight burst of exonuclease activity on the YTMA substrate, although the subsequent rates of excision were the same between the two substrates (Fig. 4B). In contrast, YktR5 was able to completely digest the YTMA substrate at the same rate as the unmodified substrate. Given that the YTMA lesion stabilizes the DNA duplex ^20^, this is consistent with YtkR5’s preference for a dsDNA substrate and suggests that exonuclease digestion by YtkR5 constitutes an alternate YTMA-DNA repair pathway to BER by YtkR2.

**Figure 4.**
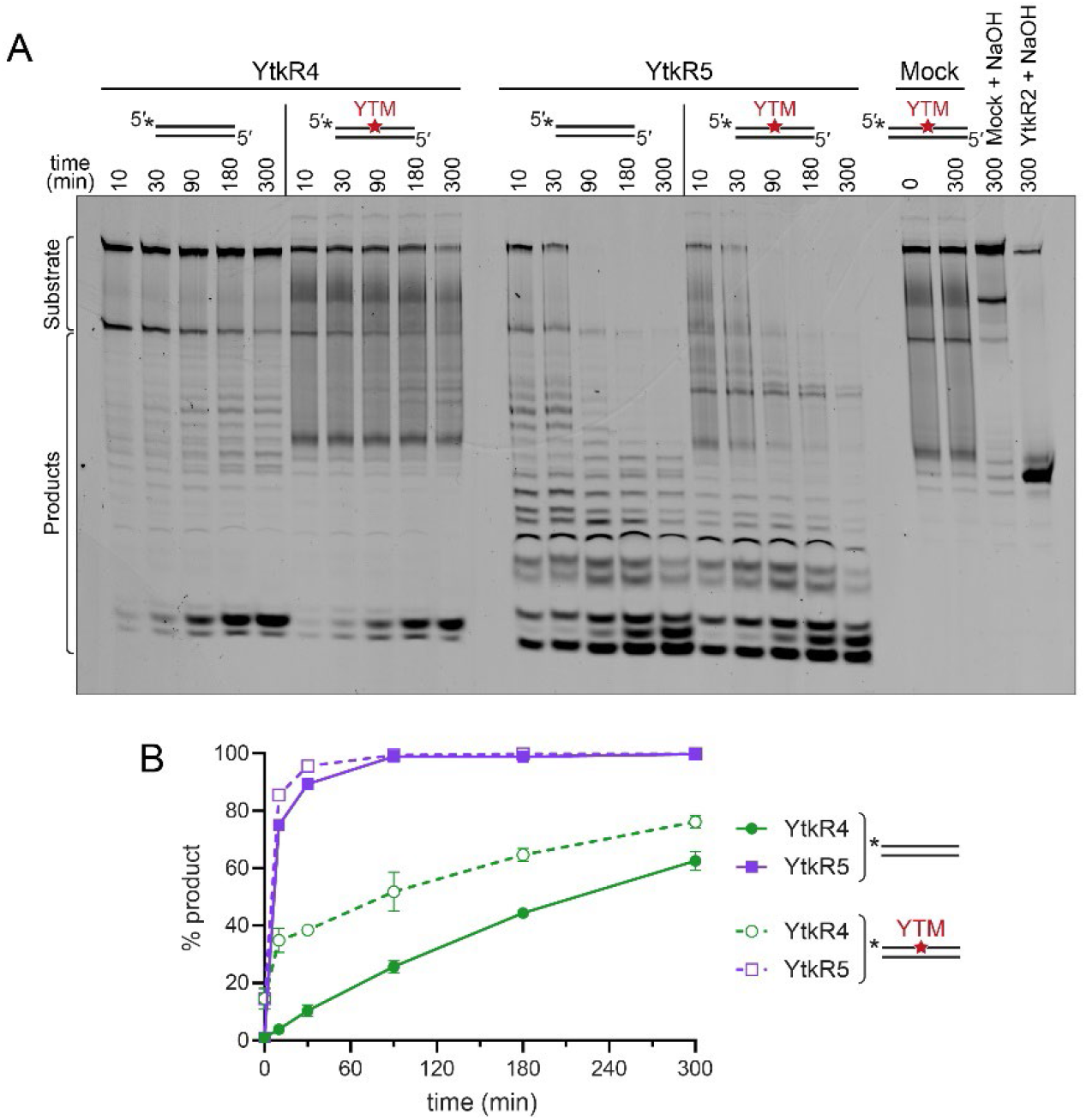
YTMA lesions do not inhibit exonuclease activities of YtkR4 or YtkR5. (A) Denaturing PAGE of YtkR4 and YtkR5 nuclease products on unmodified and YTM-modified dsDNA in the presence of 10 mM MgCl_2_. Mock, no enzyme control. The two lanes at the far right are controls to quantify the percentage of spontaneous depurination of YTMA (Mock + NaOH) and the percentage of DNA alkylated with YTM (YtkR2 + NaOH). Bands quantified as substrates and products are indicated. (B) Quantification of the gel shown in panel A. Data are represented as the mean ± SD (n = 2).

### YtkR4 and YtkR5 are found together in gene clusters

The specificity of YtkR4 and YtkR5 for the product of the YtkR2 reaction is consistent with co-evolution of these proteins. To investigate whether this is unique to the *ytk* cluster, we searched for co-existence of these proteins in other genomes. We performed a BLASTp search to identify genomes that contain a YtkR4 homolog and obtained 566 hits. We then performed two additional BLASTp searches for YtkR2 and YtkR5 homologs in these genomes, of which 311 had a YtkR2 homolog and 526 had a YtkR5 homolog. Therefore, YtkR4 homologs are only very rarely present in genomes without YtkR5 homologs, while the same is not true for YtkR2. There were incomplete genomes present in our analysis so it is possible that the fraction of YtkR4 homolog-containing genomes that also contain YtkR5 or YtkR2 homologs are underestimated but we expect that the difference between YtkR5 and YtkR2 would hold for an analysis of only complete genomes.

YtkR5 is a member of the xylose isomerase-like superfamily that contains both sugar isomerases and DNA endonucleases ^54^, so we cannot be certain that all our hits have YtkR5-like activity rather than sugar isomerase activity. To account for this, we clustered enzymes by function ^55^ using a sequence similarity network (SSN). We found that there were four major sequence clusters of YtkR5 homologs as well as several outlier sequences in clusters with fewer than three sequences (Fig. 5A). We found that YtkR2 is generally distant from YtkR4 in the genome, with only a small fraction of the homologs being within 100 kpb of each other, meaning that most YtkR2 homologs are not in the same BGC as the YtkR4 homolog (Fig. 5B). Conversely, nearly all YtkR5 homologs from the four major SSN clusters are within 10 kbp of the YtkR4 homolog, making it highly likely that they are in the same BGC. This also suggests that all the major SSN clusters have YtkR5-like activity because it is unlikely that a sugar isomerase would be so tightly associated with YtkR4 homologs. The YtkR5 homologs that were in SSN clusters with three or fewer members were roughly evenly split between being within 10 kbp and being further than 100 kbp from YtkR4, which could indicate that only some of them have YtkR5-like functions (Fig. 5). This analysis agrees with the gene co-occurrence patterns described above in which YtkR4 and YtkR5 homologs co-occur more often than the YtkR4 and YtkR2 homologs.

**Figure 5.**
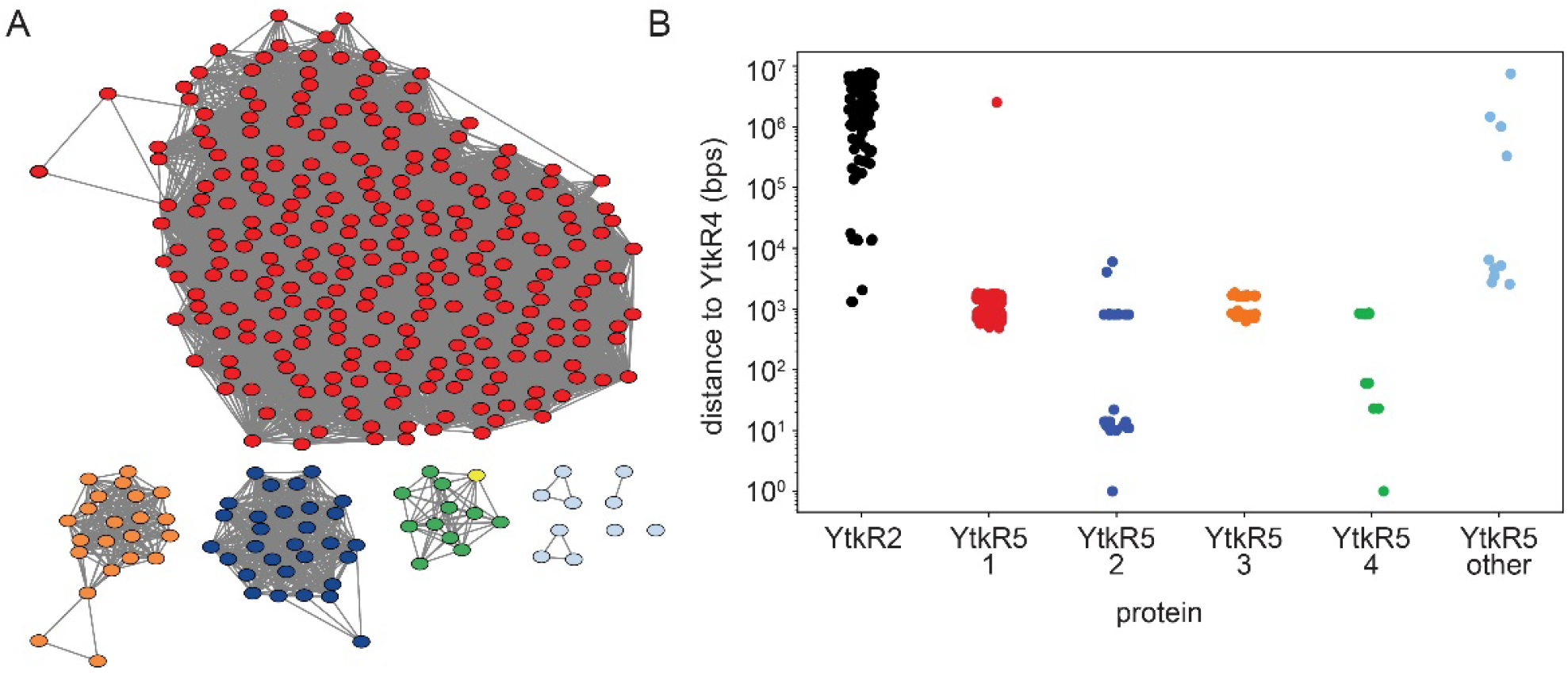
YtkR4 and YtkR5 are found together in clusters. (A) Sequence similarity network (SSN) of BLASTp hits for YtkR5. Nodes are clustered by sequence similarity and colored based on cluster membership, with all clusters with three or fewer members colored light blue. The YtkR5 query sequence is colored yellow. (B) Distance in base pairs in the genome between YtkR4 and YtkR2 or YtkR5 BLASTp hits. YtkR5 hits are further divided into subclasses based on the SSN, with colors corresponding to the SSN in panel a. The “YtkR5 other” category includes all proteins that were in a cluster with three or fewer members.

To better understand the potential non-redundant roles of YtkR4 and YtkR5, we examined the nature of the gene clusters that contained both homologs. We first used antiSMASH to determine if there was a predicted BGC within 40 kbp of YtkR4. BGCs were predicted for 113 out of the 566 YtkR4 hits. To further analyze these BGCs, we used BiG-SCAPE ^56^ to cluster both the regions identified by antiSMASH as BGCs and the regions surrounding YtkR4 hits that were not identified as BGCs. This analysis showed that the antiSMASH-identified BGCs are diverse, with the largest gene cluster family (GCF) consisting of only five members. There were two GCFs containing antiSMASH-identified BGCs with more than four members that had high similarity to known BGCs; one was made up of BGCs with high similarity to the yatakemycin BGC and the other had high similarity to the BGC that produces fluostatins M-Q (Supplementary Fig. S5). While fluostatin M-Q have not been described to have cytotoxic or antibacterial activity, structurally dimeric fluostatin compounds have antibacterial activity and these dimeric compounds have similar structures to the known DNA damaging natural product lomaiviticin A ^57^.

In contrast to the antiSMASH-identified BGCs, regions surrounding YtkR4 and YtkR5 homologs that were not identified as containing an antiSMASH BGC had larger GCFs, with the largest GCF having 20 members. Further examination of these GCFs revealed that several contained proteins with annotations similar to YtkR4 and YtkR5 homologs, specifically type I phosphodiesterase/nucleotide pyrophosphatase, an additional xylose isomerase TIM barrel, UbiA prenyltransferase, and Myo-inositol-phosphate synthase. A literature search of these domains revealed that proteins containing some of these domains are found in the *ebo* gene cluster (Supplementary Fig. S6), which is widespread among cyanobacteria and algae. In cyanobacteria this cluster has been linked to the production of the natural product scytonemin ^58^. Another variant of *ebo*, EDB, has been discovered in *Pseudomonas fluorescens* NZI7 and is responsible for the production of indole-derived compounds that repel *C. elegans* ^59^. One of the genes in the *ebo* cluster, EboB, contains a predicted TatD protein with low sequence similarity to YtkR4 (approximately 24% sequence identity). EboB’s function has not been determined, although knockouts of EboB show reduced scytonemin production ^58^. No reported *ebo* cluster from cyanobacteria contains a protein with similarity to YtkR5.

To learn more about the evolution of YtkR4 homologs and EboB and their relationship to YtkR5, we constructed a phylogenetic tree of our YtkR4 hits, EboB proteins from cyanobacteria and *P. fluorescens*, and TatD proteins (Supplementary Fig. S7). This tree revealed that YtkR4 homologs that lack a nearby YtkR5 homolog are scattered throughout the tree and are not confided to any single clade. This is also true when looking at YtkR4 homologs that have at least one nearby homolog of a gene from the *ebo* gene cluster (excluding EboB). This suggests that the ancestor of YtkR4/EboB likely cooccurred with YtkR5, and that the YtkR5 homolog was later lost in the cyanobacterial *ebo* cluster and in some other branches of the tree. Given the role of the *ebo* cluster in natural product production, it is likely that the *ebo*-like clusters containing both YtkR4 and YtkR5 homologs are involved in facilitating the production of and providing resistance to DNA-damaging natural products produced by BGCs elsewhere in the genome, possibly with lower specificity than YtkR4 and YtkR5.

Together, these results suggest that the YtkR4 and YtkR5 homologs are both important for resistance to other DNA damaging natural products and therefore often co-occur in the same genome or BGC. This co-existence is consistent with their different nuclease activities, which would provide non-redundant mechanisms for lesion repair (Fig. 6). The weak co-existence and genomic distance between YtkR2 and YtkR4 homologs in other genomes suggests that those YtkR2 homologs do not rely on a YtkR4 or YtkR5 nuclease to process the AP site product. Indeed, homologous *ytkR4* and *ytkR5* genes are not present in the CC-1065 BGC ^22, 60^. Similarly, there is a class of BGCs that contain an AlkZ-like glycosylase, unrelated to YtkR2, which provides self-resistance to DNA crosslinking and intercalating natural products ^37, 61, 62^, and there are no apparent AP endonucleases or other DNA repair genes to remove the AP lesion in those clusters. In those cases, it is unclear how the AP site products are resolved.

**Figure 6.**
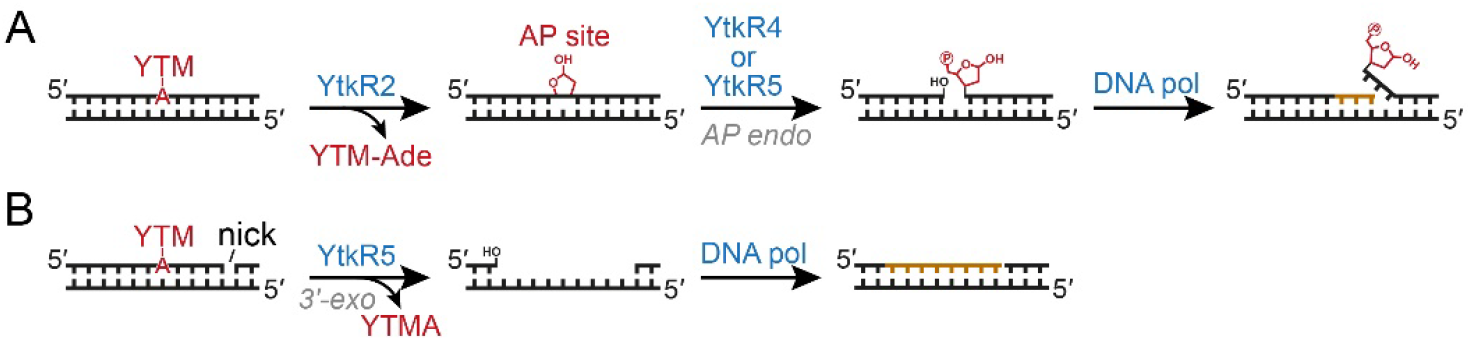
Two possible mechanisms for repair of a YTMA-DNA adduct. (A) The YtkR2 glycosylase initiates a BER pathway by excising YTM-adenine. The AP site product is incised by YtkR4 or YtkR5 to generate a 3′-OH, which is a substrate for 5′-deoxyribosephosphate displacement synthesis by a DNA polymerase. (B) The 3′-exonuclease activity of YtkR5, acting at a downstream nick in the DNA, is capable of removing the YTMA nucleotide.

In the specific case of YTM, our genomic analysis suggests that the specificity of *Streptomyces* sp. TP-A0356 YtkR4 and YtkR5 for the product of the YtkR2 reaction, and thus the existence of at least a partial BER system for this lesion, is unique to the *ytk* cluster. Our genetic deletion of YtkR4 and YtkR5 in the YTM producing strain suggests that these nucleases operate non-redundantly to enable YTM production and is consistent with their unique substrate specificities. However, the need for two separate nuclease-dependent pathways for YTMA repair is unclear. Based on the remarkable stability that a YTMA adduct imparts to the DNA double-helix ^20, 22^, it stands to reason that multiple repair pathways are needed to fully remove YTM-derived DNA adducts. Indeed, we found that both AlkD-dependent BER and UvrA-dependent NER pathways are operative in YTMA repair in *Bacillus cereus* ^20^. It may be that each nuclease repairs YTMA from a different sequence or genomic context, or that one is specific for YTMA byproducts yet to be identified. Regarding YtkR4 and YtkR5 homologs in other BGCs that lack a DNA glycosylase, we speculate that these enzymes are not AP endonucleases and that any putative exonuclease activity enables repair of a broad range of DNA adducts and toxic DNA repair intermediates. More work is needed to understand the particular repair strategies employed by these enzymes in YTM producing and non-producing bacteria.

## Experimental

### Construction and complementation of *ytkR4* and *ytkR5* mutants

Bacterial strains and plasmids used in this study are listed in Supplementary Table S1. The *ytkR4* and *ytkR5* gene in-frame deletion mutants were constructed as previously described ^60^. Briefly, two homologous DNA arms were cloned into the *Hin*dIII and *Eco*RI sites of the thermosensitive plasmid pKC1139 using primers listed in Supplementary Table S2. Then, the recombinant plasmid was introduced into *Streptomyces* sp. TP-A0356 from *E. coli* S17-1 to obtain apramycin-resistant exconjugants at 30°C. The exconjugant was then grown at 37°C to obtain apramycin-resistant single-crossover mutant culture, which was further used for screening apramycin-sensitive clones without apramycin selection for generations. The desired double-crossover mutants were verified by PCR analysis with primers listed in Supplementary Table S2.

For complementation experiments, each coding sequence was cloned into the *Nde*I and *Eco*RI sites of the integrative plasmid pIB139, and the recombinant plasmid was introduced into *ytkR4* or *ytkR5* mutant from *E. coli* S17-1 to obtain apramycin-resistant exconjugants at 30°C. The apramycin-resistant exconjugants were then used for further fermentation analysis.

### Fermentation and analysis of YTM metabolites

For *Streptomyces* sp. TP-A0356 and its derivative strains, fermentation and analyses of YTM and other relative metabolites were carried out as previously described ^38^. Briefly, *Streptomyces* sp. TP-A0356 and its derived mutant strains were first inoculated in the liquid seed medium (Tryptic Soy Broth) in a 250 mL flask, and then the culture was used for transfer onto the solid fermentation medium (International Streptomyces Project medium 2) for growth at 30°C for ∼5 days. The culture was used for extracting YTM and other metabolites, which were for further HPLC analysis.

The HPLC analysis was performed on the Agilent 1200 HPLC system (Agilent Technologies Inc., USA) using a reverse-phase Alltima C18 column (5 μm, 4.6 × 250 mm). Solvent A was H_2_O and Solvent B was CH_3_CN, and the flow rate was 1 mL min−1 with DAD detector. The analytic HPLC conditions are as follows: the gradient program was 0–3 min 15% B, 3–6 min 15–40% B, 6–12 min 40% B, 12–19 min 40–55% B, 19–22 min 55–85% B, 22–28 min 85% B, and 28–29 min 15% B.

### Protein expression and purification

The coding sequences of YtkR4 and YtkR5 were synthesized by GenScript without codon optimization and ligated into pBG102, a modified pET-27 expression vector encoding an N-terminal Rhinovirus 3C-cleavable hexahistidine-SUMO fusion tag. For YtkR4 expression, the plasmid was transformed into C41 cells along with the pG-Tf2 vector (Takara Bio) encoding Trigger Factor, GroES, and GroEL chaperones. Cells were grown at 37°C in Luria Bertani (LB) medium supplemented with 5 µg/L tetracycline. When the cultures reached an A_600_ of 0.6, they were incubated for 1 hr at 18°C before protein expression was induced by addition of 0.1 mM IPTG. Induced cells were grown overnight at 18°C. The plasmid encoding YtkR5 was co-transformed into BL21 cells along with the pGro7 vector (Takara Bio) encoding GroES and GroEL chaperones. Cells were grown at 37°C in LB medium supplemented with 5 g/L arabinose, as well as 2 mM betaine and 50 mM sorbitol to increase protein solubility. When the A_600_ reached 1.0, the cells were incubated for an hour at 18°C and 0.09 mM IPTG was added to induce the protein expression. Cultures were then incubated at 18°C overnight.

For both YtkR4 and YtkR5, cells were harvested by centrifugation and the resulting cell pellet was resuspended in lysis buffer (30 mM MOPS pH 7.5, 300 mM NaCl, 10% glycerol). Benzonase (25 U/L culture, Sigma-Aldrich) was added to the lysis buffer at a concentration of 25 U/L culture, as well as 6 mM MgCl_2_, 10 mM KCl, and 4 mM ATP to detach the chaperones from the proteins during cell lysis. Cells were lysed using an Avestin C3 Emulsiflex operating at 15,000 psi. Cell debris was removed by centrifugation at 45,000 x g for 30 min. The supernatant was supplemented with 40 mM imidazole and applied to a Ni-NTA column. The column was washed with 20 column volumes of lysis buffer supplemented with 40 mM imidazole and the protein was eluted using buffer B (lysis buffer supplemented with 500 mM imidazole). The hexahistidine-SUMO tag was removed by overnight cleavage at 4°C while dialyzing against the lysis buffer to remove imidazole. The sample was then reinjected onto the Ni-NTA column and the flow-through collected. For YtkR4 only, an additional purification step was performed, whereby the protein was then diluted four times in buffer Q (20 mM HEPES pH 7.5, 5% glycerol) and injected onto a Q-sepharose column. The protein was eluted using a 0-1 M NaCl gradient. YtkR4 and YtkR5 were concentrated using an Amicon Ultracel-10 (Merck Millipore), incubated for 15 min in 10 mM EDTA, and injected over onto a HiLoad Superdex 200 16/60 pg column (Cytiva) equilibrated in buffer S (30 mM HEPES pH 7.7, 300 mM NaCl, 5% glycerol). The proteins were concentrated to 100 µM and aliquots flash frozen in liquid nitrogen and stored at −80°C.

### DNA substrate preparation

Oligodeoxynucleotides were purchased from Integrated DNA Technologies. The sequences used are provided in Supplementary Table S3. Oligonucleotides containing a centrally located THF or deoxyuridine residue were labeled with 6-carboxyfluorescein (FAM) on the 5′-end and HPLC purified. Double-stranded substrates were formed by annealing FAM-oligonucleotides to an unlabeled complementary strand in annealing buffer (20 mM EPPS pH 8.0, 50 mM NaCl). Sequences containing natural AP sites were generated by reacting 50 µL of 50 µM dsDNA uracil-containing oligonucleotide with 5 U uracil DNA glycosylase (New England Biolabs) for 60 min at 37°C. YTM was purified as previously described ^26^ and YTM-DNA substrates were generated as previously described ^20^, except reaction mixtures contained 20 mM EPPS pH 8.0, 50 mM NaCl, 10% (v/v) dimethylsulfoxide, 10 μM DNA, and 150 μM YTM. After 18 hr at 22°C, excess YTM was removed by passing the reaction mixture through a Microspin G25 column (Cytiva) equilibrated in annealing buffer.

### Nuclease assays

Nuclease reactions were carried out at 37°C with 10 µM protein, 100 nM DNA, and buffer containing 20 mM EPPS pH 8.0, 50 mM NaCl, and 10 mM MgCl_2_. Experiments to test activity in the presence of different divalent metals were carried out under the same conditions but supplemented with either 5 mM CaCl_2_, 0.05 mM ZnCl_2_, 3 mM MnCl_2_, 1 mM NiCl_2_, 1 mM CuCl_2_, or 10 mM EDTA. Reactions were quenched by addition of an equivalent volume of loading buffer (80% formamide, 25 mM EDTA pH 8.0, 2 mg/mL orange G, and 1 mg/mL xylene cyanol, 10 U proteinase K), heated at 70°C for 10 min, and subjected to denaturing polyacrylamide gel electrophoresis (PAGE) on 20% acrylamide/8M urea sequencing gels. Fluorescence from the FAM-labeled DNA was detected using a Typhoon Trio variable mode imager (GE Healthcare). For excision of the YTM adducts, 10 µM YtkR2 was added to the reactions prior to YtkR4 or YtkR5. To verify excision of the YTM-adduct by YtkR2, reactions were quenched by addition of 100 mM NaOH and heated at 70°C for 10 min before addition of loading buffer. For quantification of nuclease activity, bands corresponding to substrates and products, as labeled in Figures 2–4, were integrated using ImageQuant (Cytiva) software.

### Identification of YtkR4, YtkR2, and YtkR5 homologs

BLAST version 2.8.1 command line software was used to perform a blastp search against the refseq_protein database with YtkR4 as the query sequence. The cutoff E-value was set at 10^-20^ and the maximum target sequences was set at 100,000. The output format was set with the following options: “7 qacc sacc sgi evalue qstart qend sstart send”. Genomes containing hits were obtained by first downloading the record for the protein from NCBI, extracting the genome accession, and then downloading the genome. Downloads were performed using the NCBI datasets command line tool using a custom script, download_blast_hit_genomes.py. A full list of genomes downloaded and used in subsequent analysis is available in Supplementary Table S4. We used tblastn to confirm presence of a YtkR4 homolog and search for YtkR2 and YtkR5 homologs in the downloaded genome. One BLAST search was run for each gene-genome pair, again using the command line with an E-value of 10^-5^ and output format options: “7 qacc sacc sgi evalue pident qstart qend sstart send”. This process was automated with a custom script, search_for_other_genes.py. The SSN was created by uploading the fasta file containing YtkR5 blastp hits to the EFI-Enzyme Similarity Tool server ^63^ and using a score cutoff of 80. The SSN was colored by cluster membership and visualized using Cytoscape.

### Analysis of genomic distance between YtkR4 and YtkR2/YtkR5

Complete genomes were determined to be circular or linear from the genbank file record using a custom script (genomes_circular_list.py). To determine the distance for a linear genome, we identified the minimum distance between the end of one gene and the start of the other. To determine the distance for a circular genome we measured the minimum number of nucleotides between genes looking in both directions in the chromosome. For genomes that were not complete, we were unable to analyze the distance in both directions. For these genomes, we assumed that if the two genes were observed on the same contig that the shorter distance between them would be their distance on the contig rather than the other possible distance if the genome is circular. Distances were not determined for any pairs that did not fall in the same contig and these pairs are not represented in the plot in Fig. 5. Distances were determined using the custom script search_for_other_genes.py. These distances were then plotted as a strip plot using the python modules seaborn and matplotlib using a custom script (make_stripplots.py). To analyze if YtkR4 was in a putative BGC, the 40 kbp on either side of YtkR4 was extracted and used as input for antiSMASH version 6 ^64^ (Supplementary Table S4).

### Construction of phylogenetic trees

TatD sequences were obtained from the Uniprot databases by searching for the term TatD and identifying the top hits with reviewed status from bacteria. EboB sequences were obtained by blasting YtkR4 against genera previously described to harbor *ebo* clusters and using the top hit. The sequences were aligned using the hmmalign program ^65^, using the TatD PFAM hmm model (PF01026) downloaded from InterPro. The phylogenetic tree was constructed using RAxML ^66^ with the following options “-m PROTGAMMAWAG -p 1234 -x 1234 -# autoFC”. We determined if EboA, EboC, EboF, or EboE hits were within 10 kb of the YtkR4 using the same methodology described above for YtkR5 using Ebo protein reference sequences determined by comparing the *ebo* cluster to the *ebo*-like clusters in *Streptomyces*. The accession numbers for the protein sequences used as references are TDC26181.1 (EboA homolog), SMKC00000000.1, locus E1265_05040 (EboC homolog), TDC26179.1 (EboF homolog), and TDC26178.1 (EboE homolog). The tree was visualized in the interactive Tree of Life ^67^, rooted at the midpoint, and labeled.

## Conclusions

We have defined the previously uncharacterized genes *ytkR4* and *ytkR5* within the YTM BGC as resistance genes important for YTM production. YtkR4 and YtkR5 are AP endonucleases acting on the product of the YtkR2 glycosylase. Both enzymes contain 3′-5′ exonuclease activity, which in YtkR5 is most pronounced on dsDNA substrates and capable of digesting through a YTMA lesion. Genes encoding YtkR4 and YtkR5 homologs in other bacteria are often located together within putative BGCs, but without a YtkR2 homolog, suggesting YtkR4 and YtkR5 play multiple, non-overlapping roles in repair of genotoxin DNA adducts.

## Author contributions

J.D.: data curation, formal analysis, investigation, visualization, writing – original draft, writing – review & editing; H.Y.: data curation, formal analysis, investigation, visualization, writing – original draft, writing – review & editing; A.S.W.: data curation, formal analysis, investigation, software, visualization, writing – original draft, writing – review & editing; G.L.T.: funding acquisition, supervision, writing – original draft, writing – review & editing; B.F.E.: conceptualization, funding acquisition, project administration, supervision, visualization, writing – original draft, writing – review & editing.

## Data availability

Data for this article, including custom scripts for genomic analysis and fasta files containing homologous sequences are available at GitHub at https://github.com/aswalker-lab/YtkR4_bioinformatics.git. Additional data supporting this article have been included as part of the Supplementary Information.

## Acknowledgements

We thank Prof. Yasuhiro Igarashi from Toyama Prefectural University, Japan for kindly providing *Streptomyces* sp. TP-A0356. This work was supported by grants from the National Science Foundation (MCB-1928918 and MCB-2341288) to B.F.E. and the National Natural Science Foundation of China (31930002) to G.-L.T.

