## Supplementary Information for "Yatakemycin biosynthesis requires two deoxyribonucleases for toxin self-resistance"

**Table S1 | Strains and plasmids used in this study**

| Strains or plasmids | Description | Source or reference |
|---|---|---|
| <u>Strains</u> | | |
| *Streptomyces* | | |
| sp. TP-A0356 | Yatakemycin producer | Ref. [1] |
| Δ*ytkR4* | *ytkR4* deletion mutant | This study |
| Δ*ytkR5* | *ytkR5* deletion mutant | This study |
| *E. coli* | | |
| DH5a | Used for routine plasmid construction | Laboratory collection |
| S17-1 | Used for *E. coli*-*Streptomyces* conjugation | Ref. [2] |
| <u>Plasmids</u> | | |
| pKC1139 | Temperature-sensitive self-replicating plasmid in *Streptomyces* | Ref. [2] |
| pIB139 | Integrative plasmid in *Streptomyces* | Ref. [3] |
| pIB-*ytkR4* | Used for the complementation of Δ*ytkR4* mutant | This study |
| pIB-*ytkR5* | Used for the complementation of Δ*ytkR5* mutant | This study |


1. Y. Igarashi, K. Futamata, T. Fujita, A. Sekine, H. Senda, H. Naoki and T. Furumai, *J. Antibiot. (Tokyo)*, 2003, **56**, 107-113.
2. T. Kieser, M. J. Bibb, M. J. Buttner, K. F. Chater and D. A. Hopwood, *John Innes Foundation, Norwich*, 2000.
3. C. J. Wilkinson, Z. A. Hughes-Thomas, C. J. Martin, I. Bohm, T. Mironenko, M. Deacon, M. Wheatcroft, G. Wirtz, J. Staunton and P. F. Leadlay, *J Mol Microbiol Biotechnol*, 2002, **4**, 417-426.

**Table S2 | Primers used in this study**

| Primer | Sequence (5′-3′) [a] | PCR product length (bp) |
|---|---|---|
| Δ*ytkR4* mutant construction | | |
| R4KO11 | CAAGCTTTCGCTCTTGAGGTACTGCTTGGC | 2098 |
| R4KO12 | CTCTAGATGTTCGGCAAGCAGTTGGAGC | |
| R4KO21 | CTCTAGACTCGTAGTCGTCGGCGTTGC | 2225 |
| R4KO22 | GGAATTCGGTTCCTGCCCGAGAAGATGC | |
| Δ*ytkR4* mutant verification | | |
| 4KOV1 | CCTGAGCCGCAGTACGGTCTT | 655 (mutant) |
| 4KOV2 | CCGCCACCACTACCACGAGA | 953 (WT) |
| Δ*ytkR4* mutant complementation | | |
| R4PCR1 | ACATATGCCGCAGCTGCCGCCTGAGG | 897 |
| R4PCR2 | AGAATTCTCACTCCTGCCGTGCTGTCGGGACG | |
| Δ*ytkR5* mutant construction | | |
| R5KO11 | CAAGCTTCTGGTTCAGTGCGTTGCTGTTGG | 2332 |
| R5KO12 | CTCTAGACCGCCACCACTACCACGAGA | |
| R5KO21 | CTCTAGAGCCAGACAGCAGGCGACGAT | 2287 |
| R5KO22 | GGAATTCCGTTGGCGAAGCGGAACAGC | |
| Δ*ytkR5* mutant verification | | |
| 5KOV1 | TCGGTGGTCAGCAGGCGTTCG | 598 (mutant) |
| 5KOV2 | AGGAAACACCGTGCTGAAGTCG | 1039 (WT) |
| Δ*ytkR5* mutant complementation | | |
| R5PCR1 | ACATATGGTGCTGAAGTCGAGCTACAACG | 837 |
| R5PCR2 | AGAATTCTCAGGCGGCAGCTGCGGCATCGT | |

[a] Underlined sequences are restriction sites

**Table S3 | Oligonucleotides used for biochemical experiments**

| Name | Sequence (5′-3′) |
|---|---|
| ssDNA | FAM-CGGGCGGCGGCA**A**A**GGGCGCGGGCC [a] |
| ssTHF | FAM-CGGGCGGCGGCAXAGGGCGCGGGCC [b] |
| ssU | FAM-CGGGCGGCGGCAdUAGGGCGCGGGCC |
| Rev | GGCCCGCGCCCTTTGCCGCCGCCCG |

[a] The nucleotides in bold indicate positions modified by YTM, with the underlined adenine being preferentially alkylated.

[b] X = THF (tetrahydrofuran)

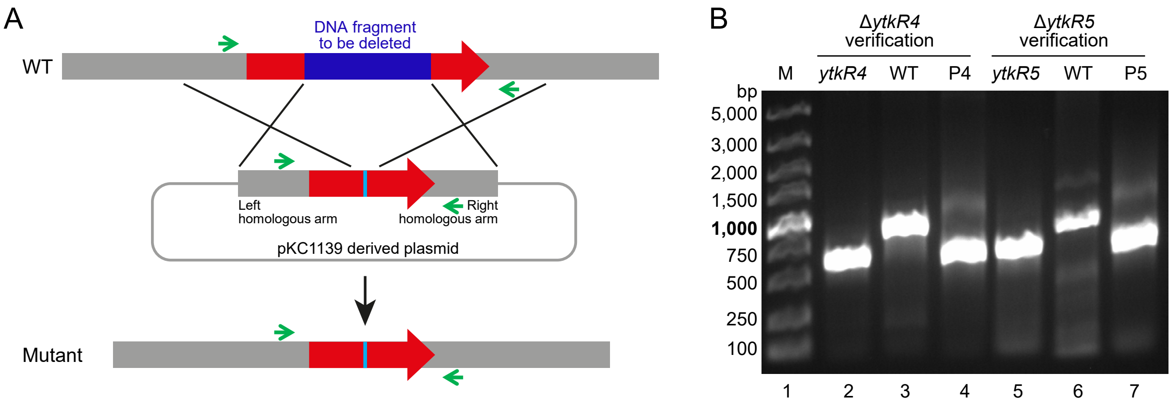


**Figure S1. Construction and verification of ΔytkR4 and ΔytkR5 mutants. A**. Schematic of the construction of the Δ*ytkR4* and Δ*ytkR5* mutants. The green arrows indicate the location of primer pair binding sites used to verify the mutants. **B**. PCR verification of the Δ*ytkR4* and Δ*ytkR5* mutants. Lane 1, molecular weight markers. Lanes 2-4, PCR products using 4KOV1/4KOV2 primers (Table S2) and either the *ytkR4* mutant genome (655 bp), the wild-type TP-A0356 genome (953 bp), or the pKC1139-derived target gene deletion plasmid (655 bp) as the template. Lanes 5-7, PCR products using 5KOV1/5KOV2 primers (Table S2) and either the *ytkR5* mutant genome (598 bp), the wild-type TP-A0356 genome (1039 bp), or the pKC1139-derived target gene deletion plasmid (598 bp) as the template.

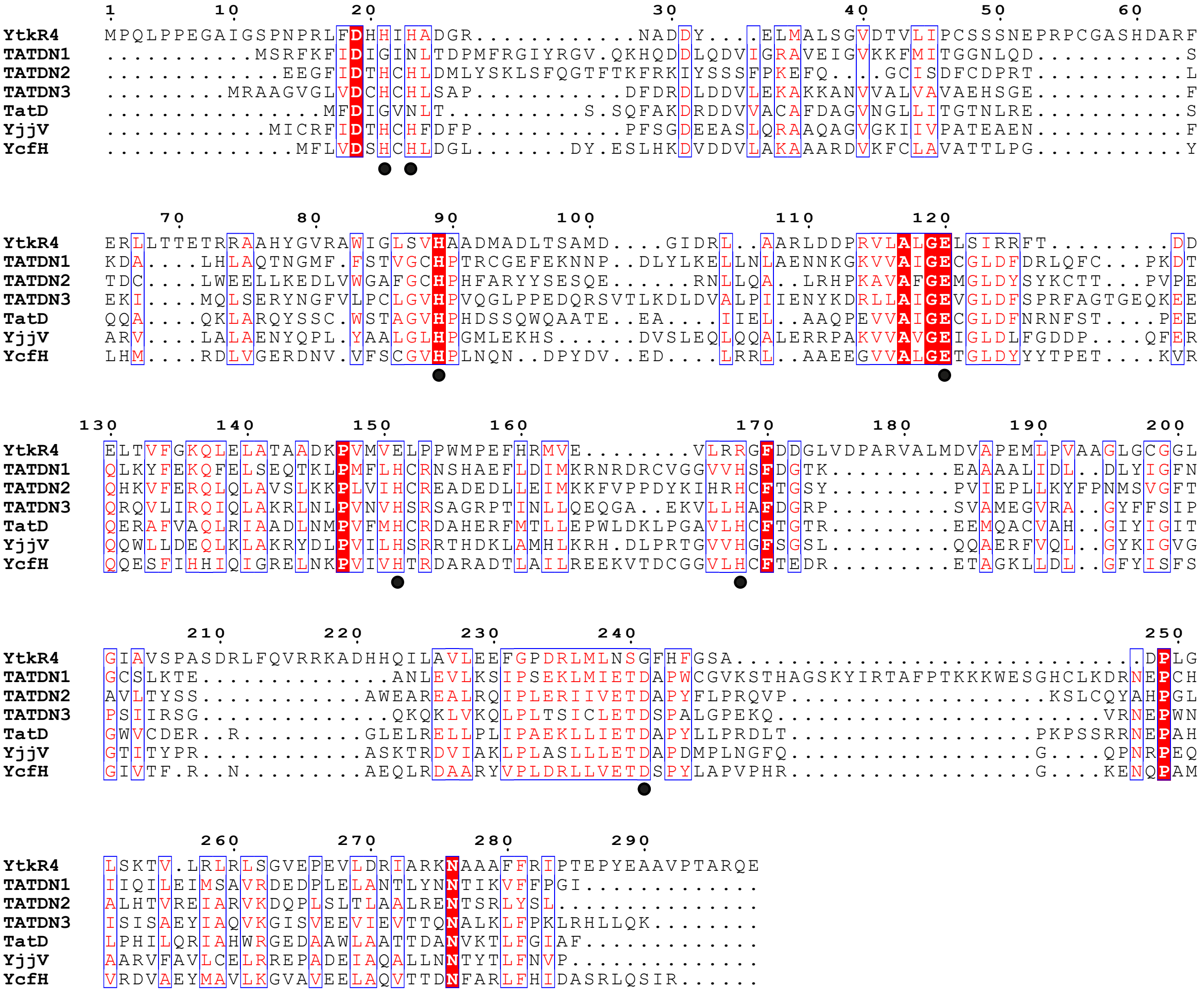

**Figure S2. Sequence alignment of YtkR4 and TatD nucleases.** Sequences are from *Streptomyces* sp. TP-A0356 (YtkR4), *Homo sapiens* (TATDN1-3), and *Escherichia coli* (TatD, YjjV, YcfH). Black circles indicate the residues involved in metal binding in TatD proteins. Sequence alignments were performed using Clustal Omega (Sievers, *et al.*, *Mol Syst Biol*, 2011, 7, 539) and illustrated with ESPript (Robert and Gouet, *Nucleic Acids Res*, 2014, 42, W320-324).

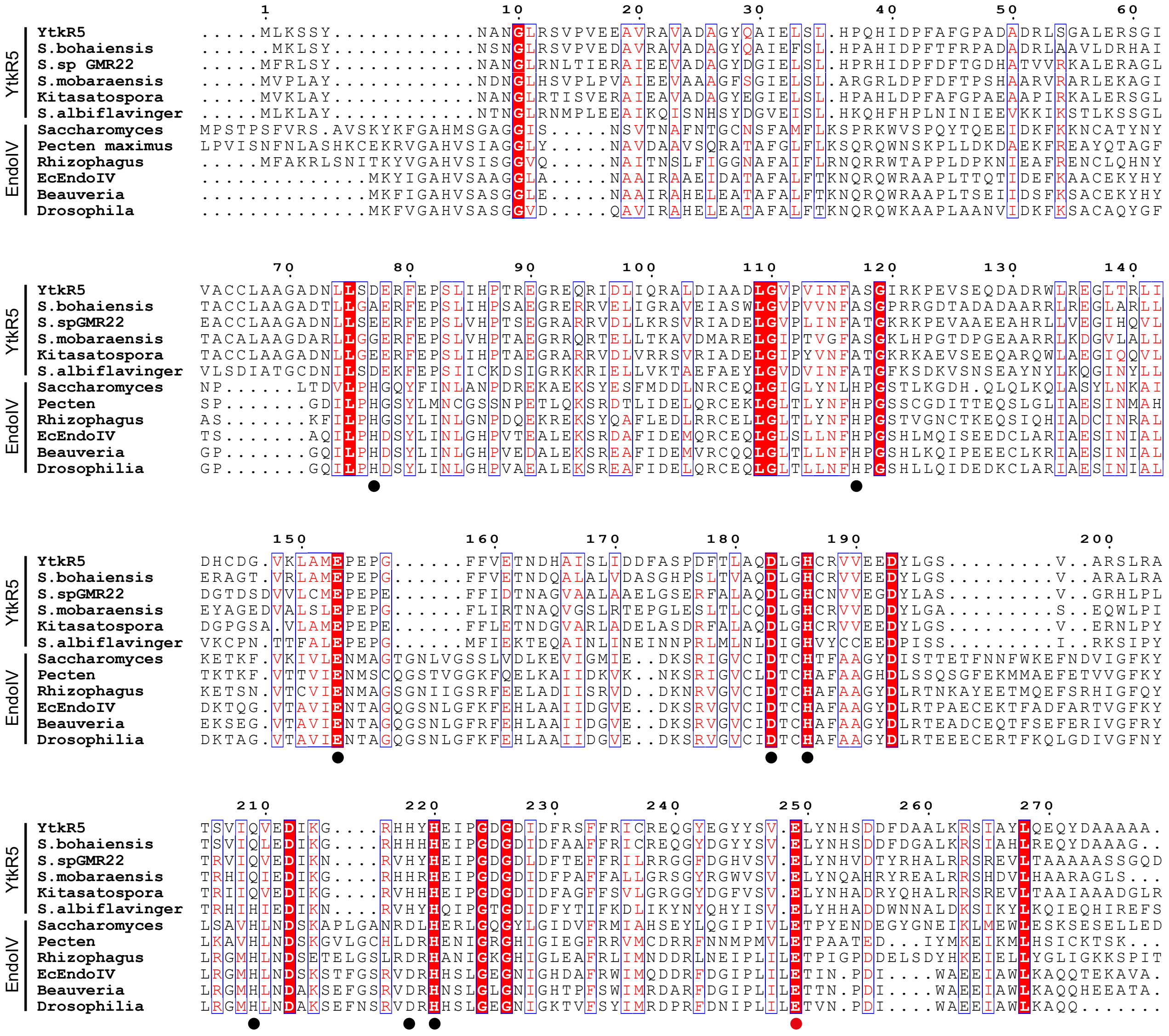

**Figure S3. Sequence alignment of YtkR5 and EndoIV homologs.** The sequence of YtkR5 was used to identify homologs using BLASTp (Altschul, *et al.*, *J Mol Biol*, 1990, 215, 403-410). Sequence alignments were performed using Clustal Omega (Sievers, *et al.*, *Mol Syst Biol*, 2011, 7, 539) and illustrated with ESPript (Robert and Gouet, *Nucleic Acids Res*, 2014, 42, W320-324). Sequences are from *Streptomyces* sp. TP-A0356 (YtkR5), *Streptomyces bohaiensis*, *Streptomyces* sp. GMR22, *Streptomyces mobaraensis*, *Kitasatospora* sp. A2-31, *Streptomyces albiflavinger*, *Saccharomyces cerevisiae* (APN1), *Pecten maximus*, *Rhizophagus irregularis*, *Escherichia coli* (EndoIV), *Beauveria bassiana*, and *Drosophila immigrans*. Circles beneath the alignment indicate the EndoIV residues important for metal binding, with the red circle denoting the catalytic Glu261.

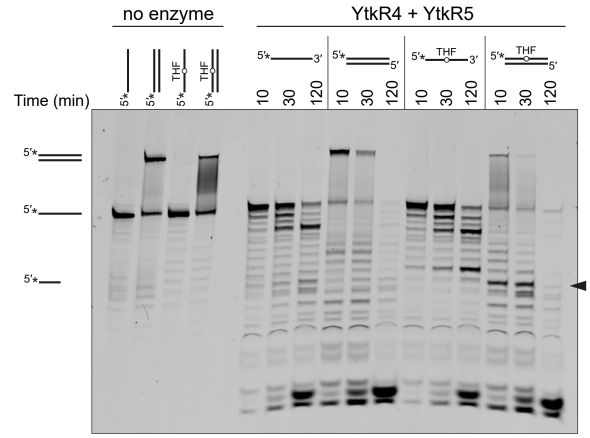


**Figure S4. Simultaneous incubation of YtkR4 and YtkR5 with DNA.** Denaturing PAGE of 5′-FAM-labeled substrates incubated with buffer (no enzyme) for 120 min or with 10 µM each of YtkR4 and YtkR5 for the indicated times. The location of the FAM label is marked with an asterisk (*). The black triangle designates the bands resulting from AP endonuclease activity. Duplex DNA persists on the gel because of the high GC content (Supplementary Table S3).

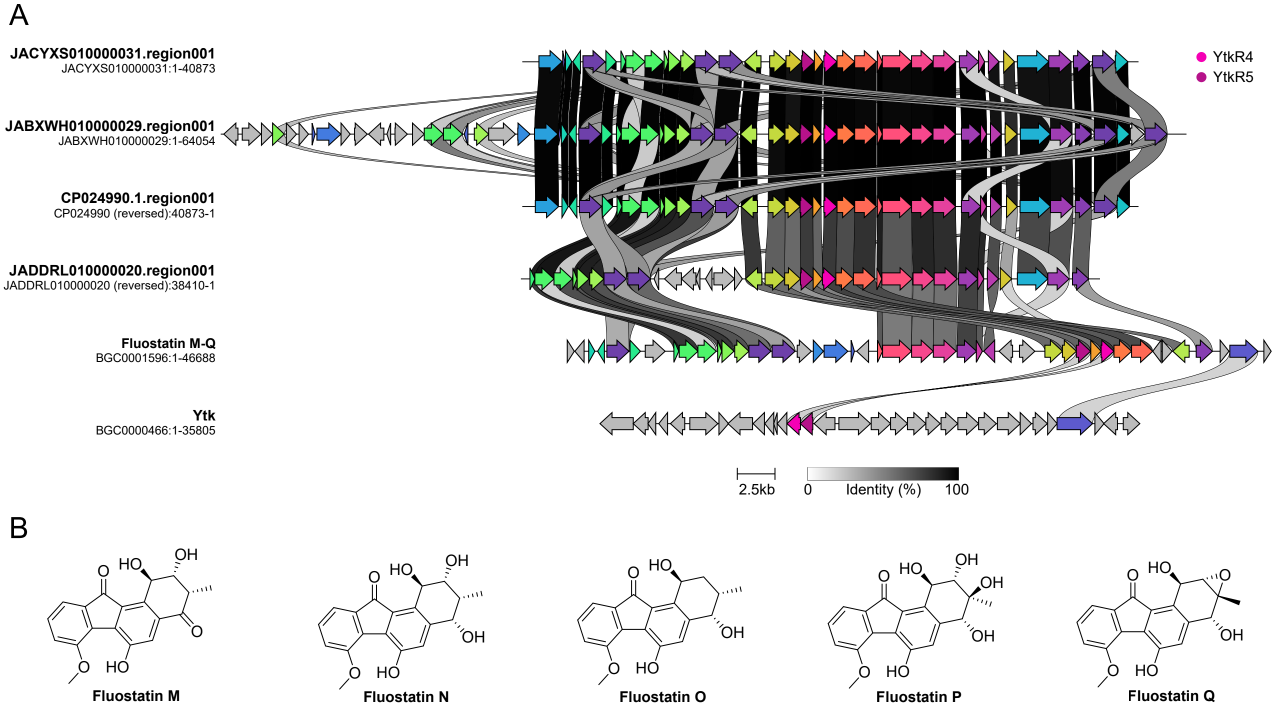


**Figure S5. Fluostatin BGCs. A**. YtkR4 hits that showed sequence similarity to the BGC for fluostatins M-Q. Fluostatin M-Q and Yatakemycin (Ytk) clusters from MIBiG [4] are shown for comparison. Genes with sequence similarity are in the same color, with YtkR4 homologs in pink and YtkR5 homologs in purple. Image made using clinker [5]. **B**. Structures of fluostatin M-Q.

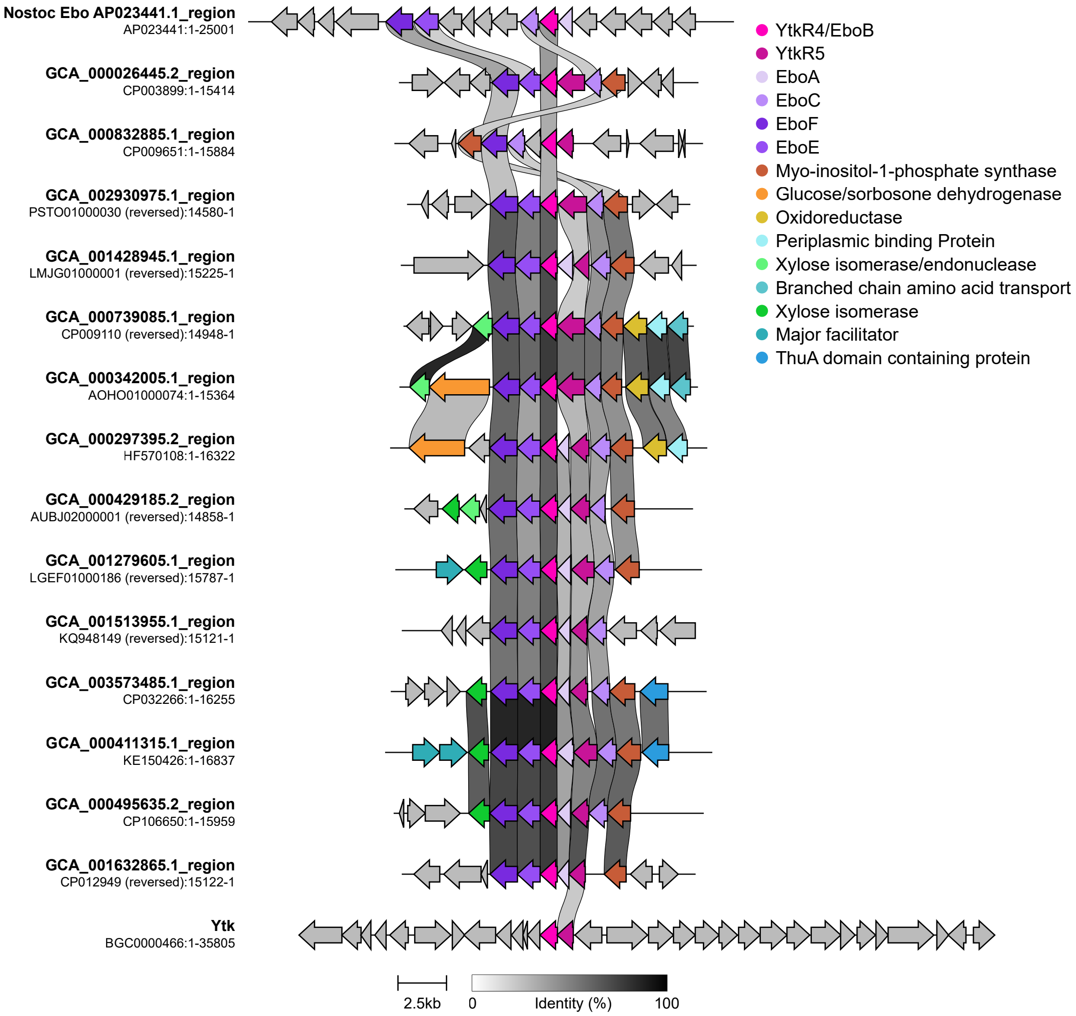


**Figure S6. Comparison of ebo-like, ebo, and Ytk clusters.** Figure shows the region surrounding the YtkR4 hits in some selected example ebo-like clusters, chosen from different BiG-SCAPE clusters. The Ytk cluster and an ebo cluster from Nostoc are included for comparison. Genes with sequence similarity are shown in the same color. Figure made with clinker [5].

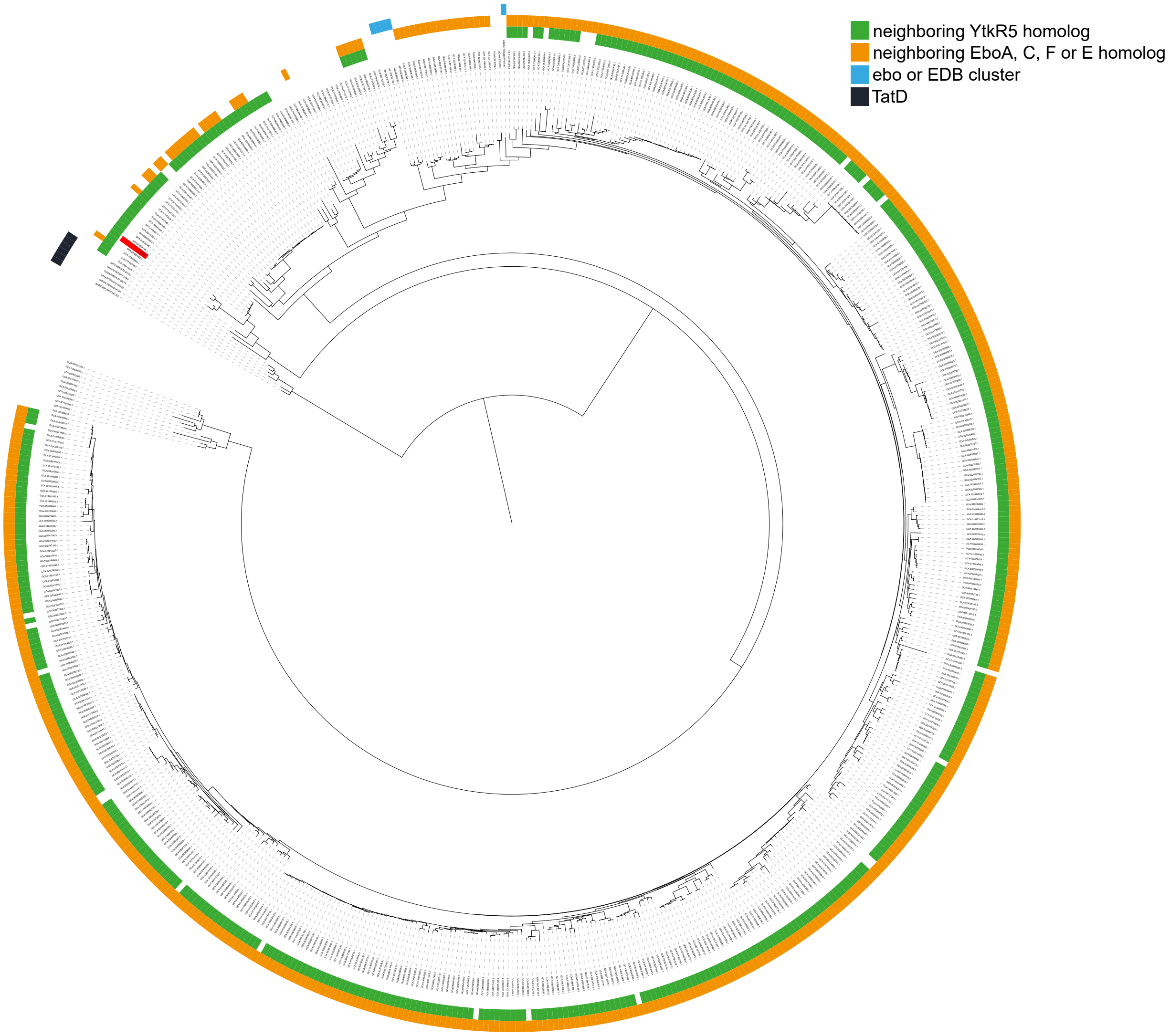


**Figure S7. Phylogenetic tree of YtkR4.** This tree shows the hits from the YtkR4 blastp search, along with bacterial TatD proteins (black squares) and EboB genes (pink squares). The initial YtkR4 query sequence is indicated by a leaf label, highlighted in red. YtkR4 homologs with a YtkR5 hit within 10 kb are indicated with a teal square. YtkR4 homologs with at least one of EboA, EboC, EboF, or EboE hits within 10 kb are indicated with a brown square. Tree and labels visualized using the interactive Tree of Life [7].
